# Gaze Direction During Bimanual Reaching Is Associated with Changes in Relative Hand Speed

**DOI:** 10.64898/2026.09.18.752785

**Authors:** Sang-Hoon Yeo, Shilokh David Sardar, R. Chris Miall, David Punt

## Abstract

The control of visually guided bimanual movements is thought to represent a compromise between the drive for synchrony that characterizes many bimanual movements and the competing demands that characterize the constituent unimanual movements. As a result, while visually guided bimanual movements are found to be ostensibly synchronized, they also demonstrate subtle asynchronies. Here, we conducted a fine-grained analysis of bimanual reaching movements focusing on their relative synchrony as movements unfold and the relationship with gaze behavior. Our novel analysis revealed that participants frequently shifted their gaze laterally several times during bimanual reaches and that these gaze shifts were systematically associated with changes in the speed ratio of the two limbs. More specifically, we observed a strong association between gaze shifts and the speed ratio shifting toward the limb on the attended side. This pattern of behavior was seen regardless of hand dominance and target difficulty, suggesting that it may reflect a common visuomotor strategy during bimanual reaching.

## INTRODUCTION

When our brain controls the movement of both hands, it prefers to control them synchronously rather than individually [1]. This bimanual synchrony appears to be a strong, prioritized control strategy, even surpassing the stringent constraints of unimanual movements such as Fitts’ law [2, 3]. However, bimanual movement is known to exhibit a subtle but systematic asynchrony alongside the overall synchrony, especially when visual guidance is crucial [2, 4–6]. It is thought that such asynchrony arises from the fact that, although both hands can move simultaneously, our gaze—that is, our overt attention—cannot be divided. For this reason, the visual guidance of bimanual movements essentially involves the brain sequentially allocating gaze to one hand or the other and modulating the kinematics of each hand in turn, resulting in asynchrony. Previous research on bimanual movement has demonstrated that overt attention to a particular limb during bimanual tasks is associated with a small but significant lead for the attended limb [7, 8].

Studies that have measured gaze patterns during bimanual reaching to a pair of targets consistently report that gaze often shifts from side to side multiple times [5, 9, 10]. The relationship between these gaze shifts and the bimanual asynchrony appears to be clear. In studies where participants are forced to fixate on one specific target or the central point between targets, there is typically greater synchrony between the limbs than in the free vision condition, although precision is diminished [5, 11, 12]. Consistent with these observations, our previous analysis of the present dataset demonstrated that visually guided bimanual reaching is characterized by strong overall synchrony together with subtle but systematic between-limb asynchronies, particularly when the two hands move toward targets with different accuracy demands [13].

Nevertheless, our understanding of the principles underlying the observed gaze-shifting patterns remains elusive. Existing analyses primarily focus on either evaluating the overall eye-movement statistics or examining snapshots of the gaze behavior, such as the initial or final gaze direction. Meaningful attempts have been made to identify different gaze strategies during bimanual reaching and relate them to movement asynchrony [4, 9, 10]. However, these analyses generally rely on task-specific classifications based on hand kinematics or movement phases defined using ad hoc criteria, making it difficult to develop a unified account of how gaze behavior relates to the evolving coordination between the two hands throughout the movement. Therefore, they provide only limited insight into how eye-hand coordination shapes the subtle asynchrony that develops throughout bimanual reaching.

Here we introduce a new analytical framework for examining eye-hand coordination during three-dimensional, unrestrained bimanual reaching movements toward a pair of targets.

Specifically, our analysis examines the ratio between left-and right-hand speeds, and how this ratio changes when gaze shifts from one side to another. By analyzing experimental eye-hand movement data acquired from healthy participants making bimanual reaching movements to a pair of targets of different sizes, we demonstrate a robust association between gaze shifts and changes in the speed ratio between the two hands.

## RESULTS

### Summary of the experiment

Participants were seated in front of a table and performed bimanual reaching movements toward a pair of circular targets displayed on a touchscreen monitor standing on the table. Initially, they placed their index fingers on the designated starting points marked on the table, while their eyes fixated on a fixation cross displayed at the center of the screen. At the beginning of each trial, the fixation cross disappeared, and circular targets on the left and right sides of the screen were presented, with their centers located approximately 39 cm away from each finger. Participants then initiated three-dimensional bimanual movements to touch the circles with their index fingers as fast and as accurately as possible. A combination of two different target sizes, small and large, was used to modulate the accuracy demands of reaching, resulting in four different bimanual reaching conditions. The positions of participants’ fingertips and their horizontal eye movements were monitored using an optical motion capture system and electrooculography (EOG). Once both fingers touched the targets, the trial ended, and participants were asked to return to the home position and wait for the next trial, while the points of touch remained displayed on the screen as visual feedback.

A total of 19 participants (10 right-handed and 9 left-handed) took part in the study, each completing 40 trials of bimanual reaching movements. This experimental protocol has been described previously [13], which focused on conventional kinematic measures such as movement timing, endpoint synchrony, and target accuracy. The present study uses the same dataset but applies a new analytical framework to examine how changes in gaze direction are associated with the evolving relative dynamics of the two hands during movement. Further details can be found in the Methods section and in Sardar et al. [13].

### Bimanual movements are largely synchronous but exhibit subtle asynchrony

To provide context for the new analyses presented below, we briefly summarize the principal findings of our previous event-based analysis of this dataset [13]. These results are included solely to establish the previously reported characteristics of the dataset and are not intended as new findings. The previous analyses focused on discrete kinematic events, including movement onset, peak speed, and movement end, together with conventional endpoint measures such as movement timing, endpoint synchrony, and target accuracy. Temporal coupling between the limbs was strong at movement onset and around peak speed, with mean signed inter-limb lags remaining within 12 ms of perfect synchrony across all bimanual conditions.

Endpoint analyses showed that the mean absolute lag increased from 67 ms during movements towards same-sized targets (congruent targets) to 106 ms during movements towards differently sized targets (incongruent targets), with the hand reaching toward the smaller, more demanding target showing a reliable tendency to arrive first. Visual orienting was likewise biased toward the smaller target regardless of whether it was presented on the dominant or non-dominant side. These event-based analyses established that endpoint asynchrony was systematically related to target difficulty and visual orienting but did not address how these differences developed continuously throughout the movement. The present study builds on those findings by examining the evolving relative dynamics of the two hands across successive gaze phases.

Although the present study does not repeat the full conventional kinematic analysis, Figure 1 provides a descriptive overview of the broad left-right coupling observed in the bimanual reaches. Across all participants and target conditions, representative kinematic variables from the two hands were strongly correlated, indicating that the movements were largely synchronous overall. This broad synchrony is important because it shows that the gaze-phase analysis below concerns subtle changes in relative hand dynamics within an otherwise tightly coupled bimanual movement.

**Figure 1.**
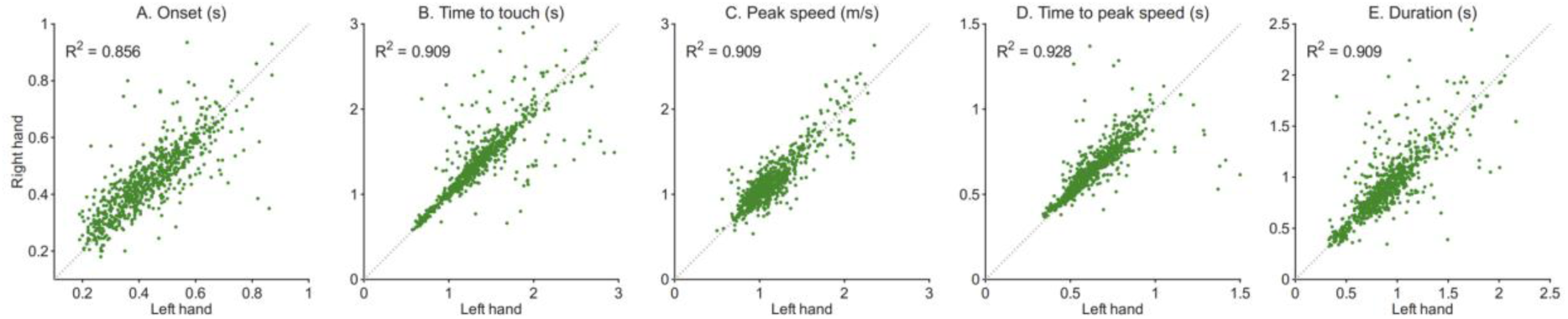
Statistics of representative kinematic variables of bimanual reaching. The horizontal axes represent values taken from the left-hand movement, while the vertical axes represent values from the right-hand. The diagonal line indicates perfect synchrony between the left-and right-hand kinematics, with the R^2^ values in the top left corner summarizing the fit to the diagonal line.

### Gaze behavior during bimanual reaching is bilateral

Figure 2 provides a descriptive overview of horizontal gaze behavior before and during bimanual reaching. During the initial fixation period, EOG signals were centered around baseline (Figure 2, gray), reflecting fixation near the central cross. After target onset, gaze was distributed predominantly toward the left (orange) or right (blue) side rather than remaining near the midline, indicating that participants typically directed overt attention towards one side or the other during the reach. This bilateral pattern of gaze behavior motivates the subsequent analysis, in which bimanual movements were segmented into left-and rightward gaze phases.

**Figure 2.**
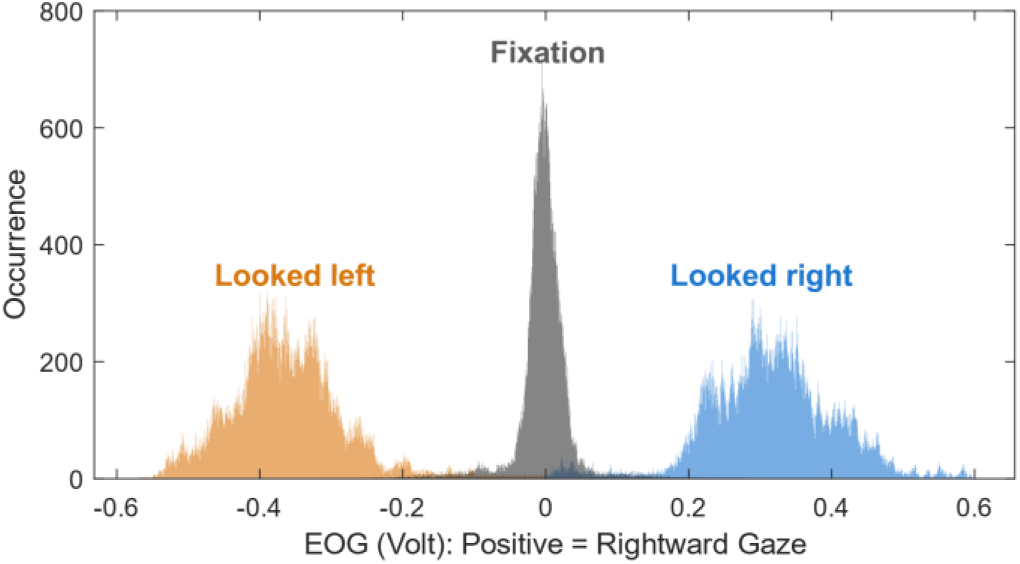
Horizontal gaze behavior before and during bimanual reaching. Histograms of EOG signals of all trials of a representative participant. Positive voltages represent the rightward gaze, and negative values represent the leftward gaze. EOG signals before the trial, when the participant was fixating on the central cross, are shown in gray, and signals after the start of the trial, when two targets were displayed, are shown in blue and orange.

### Gaze shifts during movement

Lateral gaze shifts occurred in most trials (97.3%); only 20 trials (2.7%) contained no detected lateral gaze phases. Including the initial shift away from central fixation, the number of gaze shifts was defined as the number of lateralized gaze phases within a trial. Participants made a mean of 2.20 ± 0.89 gaze shifts per trial, with a median of 2 shifts. One shift occurred in 119 trials (15.8%), whereas more than one shift occurred in 612 trials (81.5%). These results indicate that participants typically directed gaze to both sides within a single bimanual reach.

The number of gaze shifts was then examined across the four bimanual target-size combinations, with trials containing no lateralized gaze shift retained as zero-count observations. Descriptively, gaze-shift number was highest when both targets were small. A trial-level Poisson mixed-effects model with target-size combination as a fixed factor and participant as a random intercept showed a significant effect of target-size combination, *χ*^2^(3) = 11.06, *p* =.011. Model-estimated counts with the participant random effect set to zero were 2.45 gaze shifts per trial for the small-small condition, compared with 2.00, 2.05, and 2.07 for the large-large, small-large, and large-small conditions, respectively. Bonferroni-adjusted comparisons showed that the small-small condition involved significantly more gaze shifts than the large-large (*p* =.011), small-large (*p* =.028), and large-small (*p* =.045) conditions. Thus, gaze shifting increased when both targets imposed high accuracy demands, although bilateral gaze shifting was observed across all target conditions.

Figure 3 illustrates the observed gaze-shift patterns of six representative participants, with gaze direction shown across trials and aligned to the time of final target touch. In each colored matrix visualizing trials from a specific target condition for an individual participant, each column represents one trial, and time progresses from bottom to top. The pre-trial period is shown in white, the pre-movement period is shown in light gray, and the interval after movement onset but before gaze was first classified as leftward or rightward is shown in dark gray. Leftward and rightward gaze phases during movement are shown in orange and blue, respectively. Although gaze could shift during the pre-movement period, the present analysis focused only on gaze phases during the movement; therefore, pre-movement gaze shifts were not classified as leftward or rightward in this visualization.

**Figure 3.**
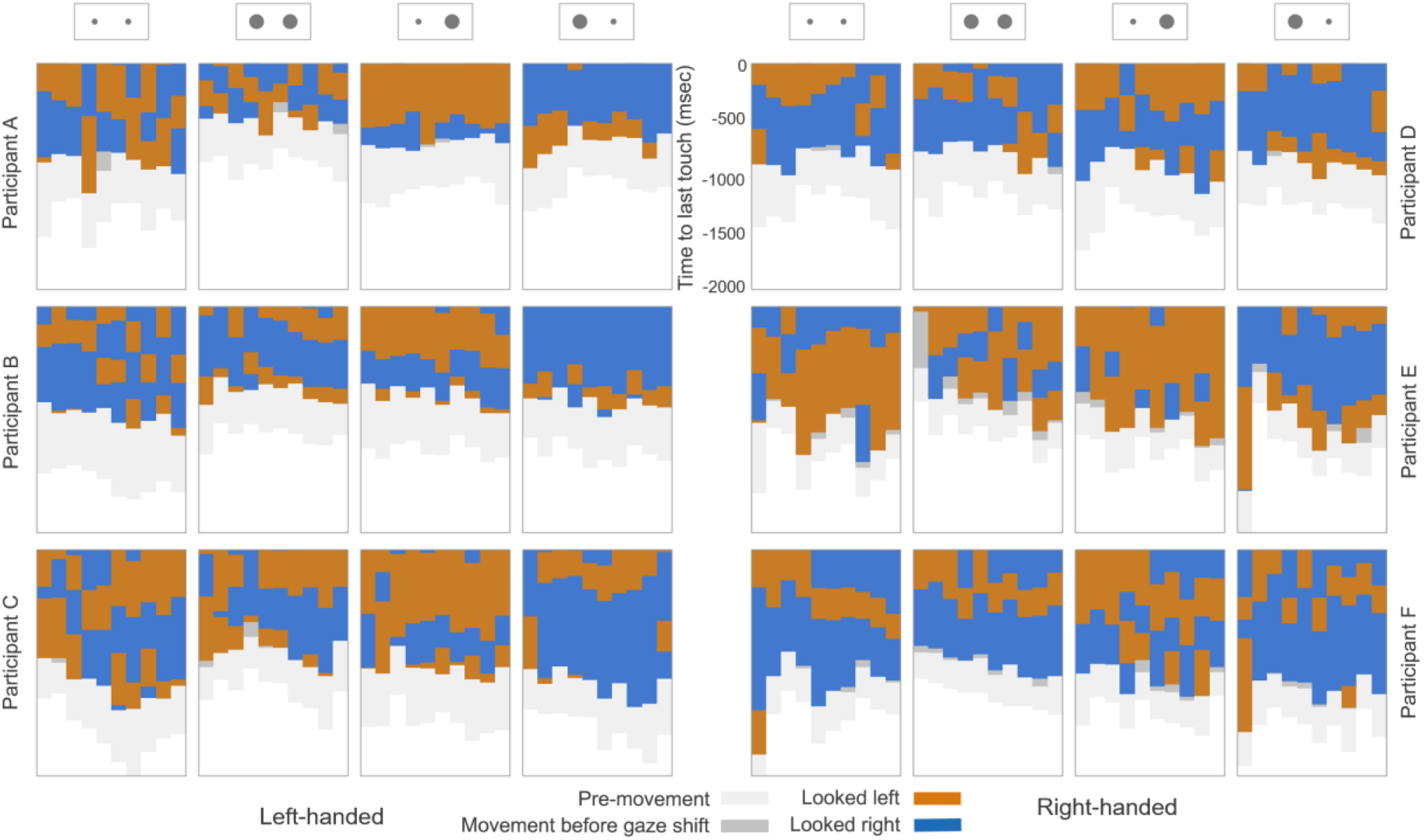
Gaze behavior of six representative participants, including three left-handed participants (A–C) and three right-handed participants (D–F), is shown as color-coded matrices. Each matrix represents trials from one target condition, with each column corresponding to a single trial. Time progresses from bottom to top and is aligned to the time of final target touch. Light gray indicates the pre-movement period, i.e., before hand movement onset and before gaze shift, while dark gray indicates the period after hand movement onset and before gaze shift. Orange and blue indicate leftward and rightward gaze phases during movement, respectively. Because movement durations varied across trials, columns differ in the amount of time represented before final target touch.

Inspection of these gaze-pattern matrices revealed substantial trial-by-trial variability. Nevertheless, the overall horizontally striated patterns within each matrix indicate that participants often used repeatable gaze-shifting patterns within a given condition. Apart from the tendency for more gaze shifts when both targets were small, there was no obvious gaze-shifting pattern shared across participants or specific to hand dominance. Therefore, although gaze behavior was not random, gaze-shifting patterns during bimanual reaching were highly variable across participants and target conditions, consistent with previous reports [2, 5]. This variability makes it difficult to identify a common strategy from gaze behavior alone and motivates the subsequent analysis of how gaze phases were coordinated with the unfolding relative dynamics of the two hands.

### Trajectory-level bimanual asynchrony is associated with gaze-shifting patterns

Here, we use the term “trajectory-level bimanual asynchrony” to refer not only to differences in final target-contact time, but more broadly to moment-to-moment differences in the relative progression of the two hands during the reach.

Examining the gaze trajectory and hand speed profiles together reveals a subtle trend in how hand speed changes when gaze shifts: the side where gaze was directed, hereafter referred to as the “seen side,” tended to show relatively greater speed progression compared to the “unseen side.” Figure 4 illustrates four representative trials of different target conditions, taken from a single participant’s data. Each figure shows the gaze trajectory with speed profiles of the left and right hand, all aligned with the time axis progressing from bottom to top. Observing the coupled dynamics between gaze and the two hands, the seen hand appears to show relatively greater speed progression compared to the unseen hand during corresponding gaze phases. For example, this trend is most noticeable in the second plot of Figure 4, where the left hand accelerated earlier and maintained higher speed when gaze was directed to the left, followed by the right hand maintaining a higher speed (i.e. less deceleration) during the deceleration phase when gaze shifted to the right. This visual examination led us to hypothesize that gaze shifts are systematically associated with the ongoing competition between the two hands, with the hand on the seen side demonstrating relatively greater increase in speed compared to the unseen hand.

**Figure 4.**
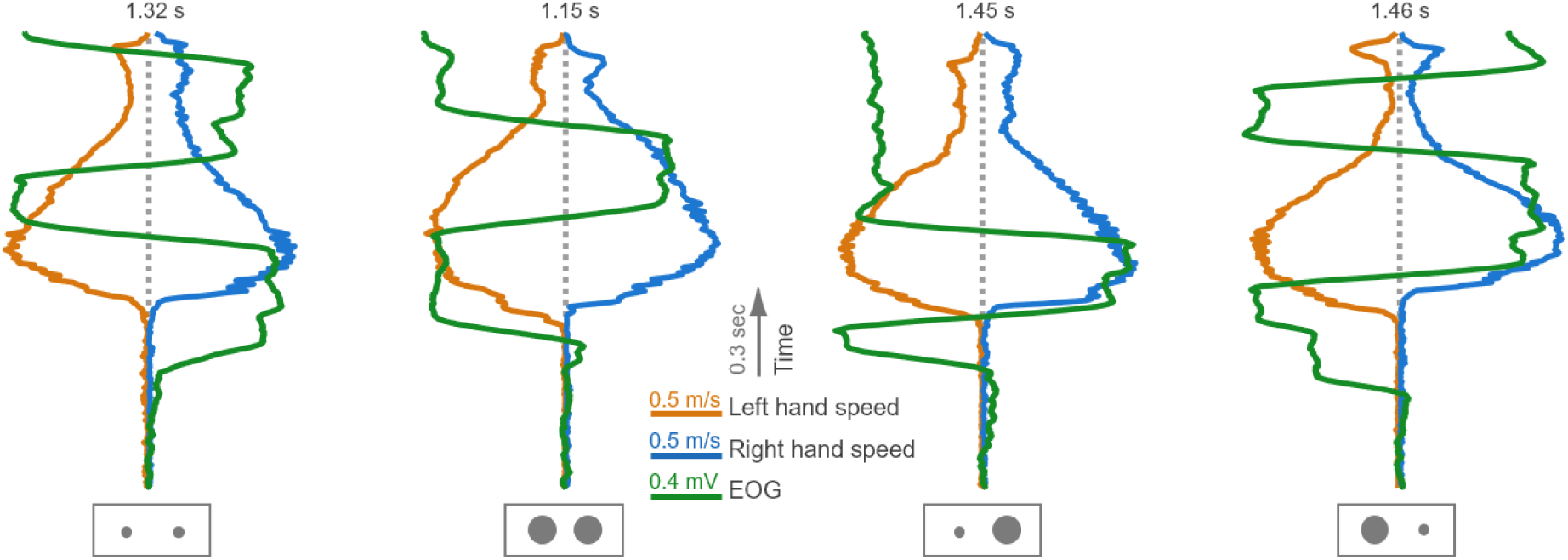
Gaze shift and bimanual speed profile. The bilateral gaze-shifting patterns (EOG profile in green) of a representative trial for each condition are plotted in synchrony with the speed profiles of both hands. The speed profiles of the left (orange) and right (blue) hands are mirrored with respect to the vertical time axis (dotted gray line), which progresses from bottom to top.

Based on this observation, our main analysis focused on investigating the competitive dynamics of the left and right hands and how these dynamics covary with gaze behavior. To highlight the competitive bimanual dynamics, the three-dimensional movement of each hand was simplified to its “traveled distance,” a one-dimensional, monotonically increasing property whose time derivative is speed. Then, in a two-dimensional space where the horizontal axis represents the traveled distance of the left hand and the vertical axis represents the traveled distance of the right hand, the competitive behavior of the two hands can be depicted as a two-dimensional trajectory, named Bilateral Traveled Distance (BTD), as shown in Figure 5 (left). Since bimanual movements are largely synchronous, BTDs are mostly aligned along the 45-degree equal-distance line, with subtle deviations from this line indicating trajectory-level asynchrony.

**Figure 5.**
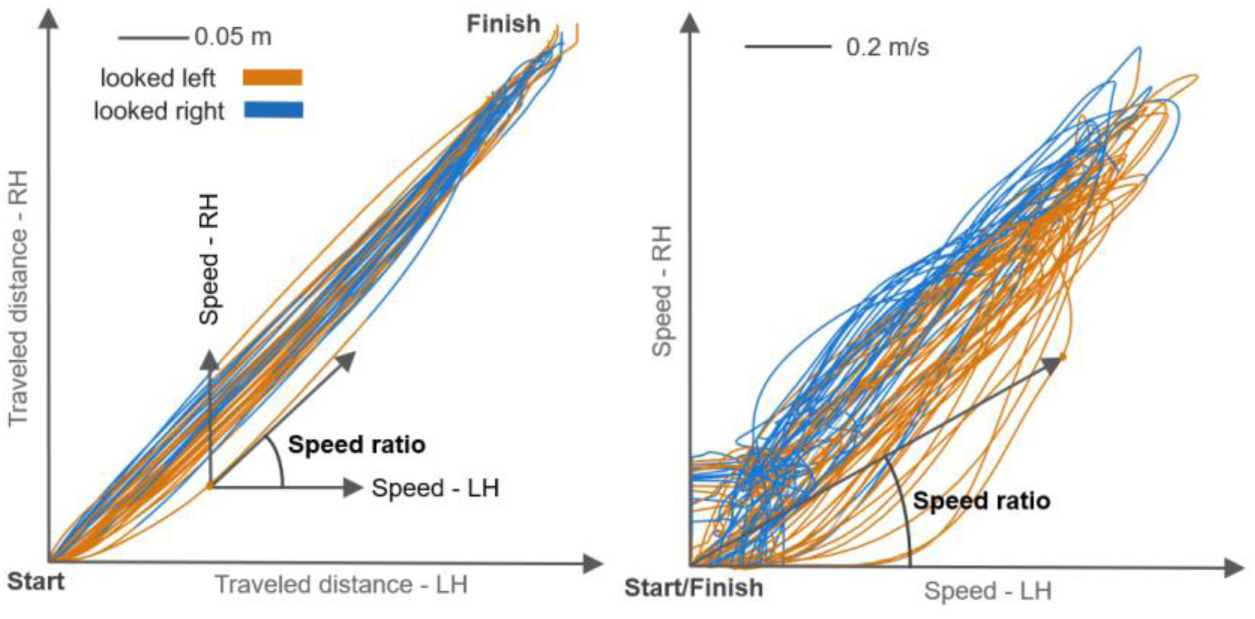
Bimanual hand trajectories in the space of traveled distance (left) and speed (right). Six-dimensional bimanual hand trajectories are represented as two-dimensional trajectories in the space of traveled distance and speed respectively. These trajectories are taken from all trials of a representative participant. In the space of traveled distances (left), bimanual reaching is depicted as a monotonically increasing curve from start (0,0) to finish. In the speed space (right), it forms a closed-loop trajectory starting and ending at (0,0). Each trajectory is segmented by gaze phase, color-coded based on looking left (orange) and looking right (blue).

At each point on the BTD trajectory, the slope represents the ratio between left-and right-hand speeds. A slope of 1 indicates that both hands are moving at the same speed; a slope less than 1 indicates that the left hand is moving faster, while a slope greater than 1 indicates that the right hand is faster. An example vectorial representation is illustrated in Figure 5 (left). This can also be visualized as the angle of a trajectory point in the space of bimanual hand speeds, as shown in Figure 5 (right). Based on this setup, the main idea of the subsequent analysis is to evaluate how gaze shifts are associated with changes in the speed ratio of the two hands, i.e., how gaze shifts and changes in BTD slope are related.

We first segmented and color-coded a BTD trajectory into discrete “gaze phases” based on gaze direction, i.e., left (orange) or right (blue), as shown in Figure 5. Afterward, the average BTD slope was calculated for each segment (see Figure 6, top-left). As bimanual movements proceed from one gaze phase to another, the average slope, i.e., the speed ratio, either increases or decreases. Given that a greater slope means a higher speed of the right hand compared to the left hand, and vice versa, an increase in slope indicates that the speed ratio shifted toward the right hand during the transition to the next gaze phase, while a decrease indicates that the speed ratio shifted toward the left hand. For convenience, we define a new variable δ, which is the difference in average slope across two consecutive gaze phases, and consider only the sign of δ.

**Figure 6.**
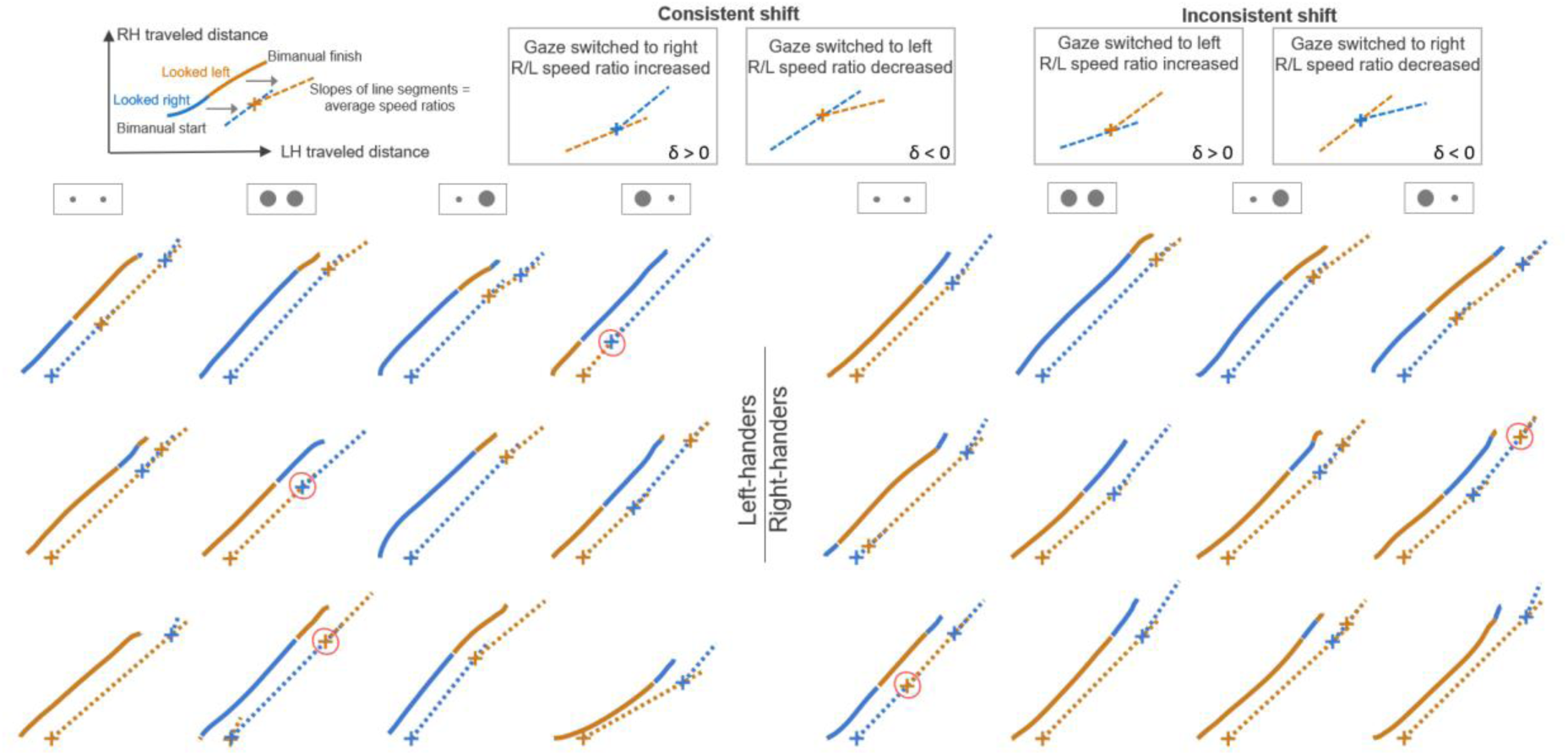
Examples of consistency analysis. A total of twenty-four trials, three for each target condition (as illustrated in the second row) and handedness (two), were randomly selected across all participants. The illustration at the top-left summarizes how average speed ratios are calculated per gaze phase: the BTD trajectory on the left is an example of two consecutive gaze phases, where each gaze phase is color-coded based on whether the gaze was on the right-(blue) or left-hand (orange) side. The dotted lines on the right represent the average BTD slopes, and the cross indicates the moment the gaze shift occurred. The dotted line of the previous gaze phase is extended beyond the cross to aid in the visual comparison of slopes. Four boxes at the top illustrate the way consistency is assessed by matching δ, the difference in average slope across gaze phases, with the gaze-shifting pattern. For each trial, a bimanual trajectory in traveled distance space with color-coded gaze phases is plotted, with corresponding average speed ratios plotted as color-coded line segments. While most shifts are consistent, inconsistent shifts are marked as red ellipses.

Based on this setup, the analysis focused on identifying how δ was associated with the gaze-shifting pattern. If gaze shifted from right to left and the slope decreased (δ negative), it would indicate that the speed ratio shifted toward the left hand, i.e., the seen side.

Conversely, if the slope increased (δ positive) when gaze shifted from right to left, it would indicate that the speed ratio shifted toward the right hand, i.e., the unseen side. The same logic was applied to shifts from left to right.

When the speed ratio shifted toward the hand on the seen side, this was termed a “consistent shift.” When the speed ratio shifted toward the hand on the unseen side, this was termed an “inconsistent shift.” All four possible cases of consistent and inconsistent shifts are summarized in Figure 6, shown in the four boxes at the top. This framework allows examination of the relationship between gaze shifts and the competitive dynamics of bimanual reaching by observing how frequently consistent shifts occur compared to inconsistent shifts.

Figure 6 shows example trials, evenly sampled from all participants and conditions, along with the corresponding consistency analysis. These examples indicate that the majority of gaze shifts and the corresponding changes in BTD slopes were consistent, with only a few exceptions highlighted by red ellipses in Figure 6.

Figure 7 summarizes the analysis results across all participants. Stacked horizontal bars represent participants, with right-handers in the upper panels and left-handers in the lower panels. Each horizontal bar represents the proportion of consistent and inconsistent gaze shifts for left (orange) and right (blue) gaze shifts. See the figure caption for a detailed interpretation of the plot.

**Figure 7.**
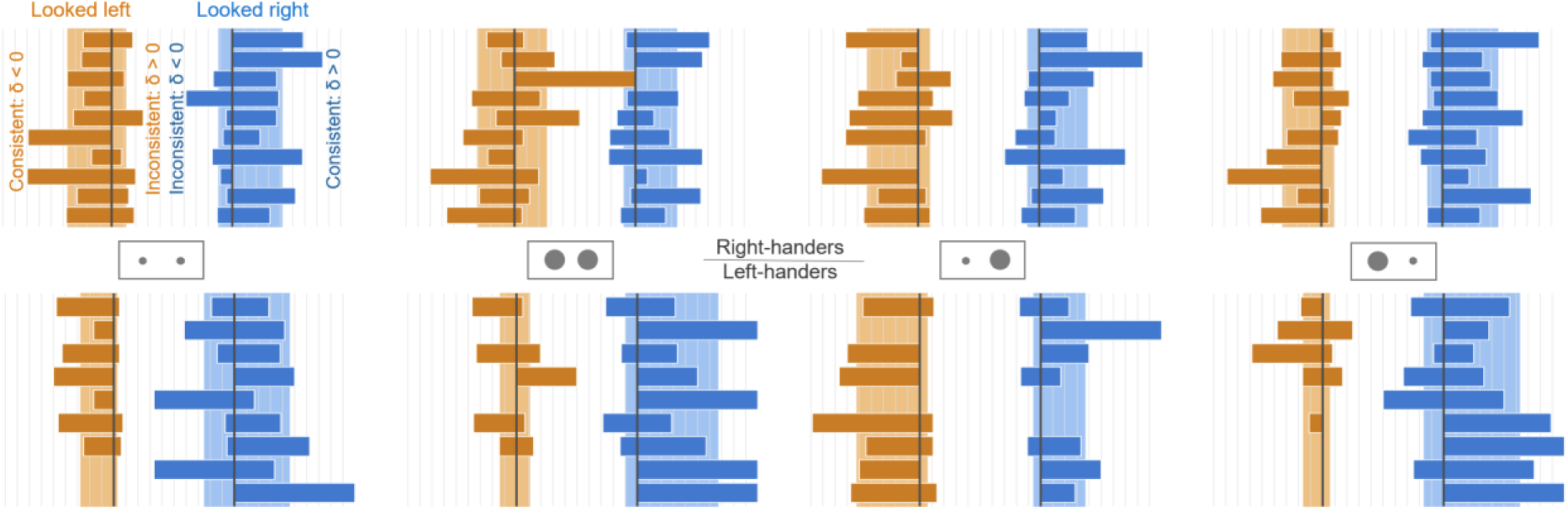
Statistics of consistent and inconsistent shifts for all participants and conditions. The upper and lower rows represent data for right-handers and left-handers, respectively. Each row consists of four pairs of vertically stacked orange/blue bars, each pair representing one of four target conditions (illustrated between the upper and lower rows). Each stacked bar represents the statistics of consistent and inconsistent gaze shifts when the participant looked left (orange) and right (blue), with each layer of the stack representing an individual participant. For the orange mirrored bars, those on the left side of the vertical line represent the proportion of consistent shifts with respect to all gaze shifts made by that participant in that target condition, whereas those on the right represent inconsistent shifts. This relationship is horizontally flipped for the blue mirrored bars, i.e., right for consistent shifts and left for inconsistent. In other words, if all orange bars are on the left of the vertical line and all blue bars are on the right, it indicates that all gaze shifts were consistent. Lighter-colored tall bars in the background of the stack represent the group average. Grid lines in the background are drawn every 10%.

**Figure 8.**
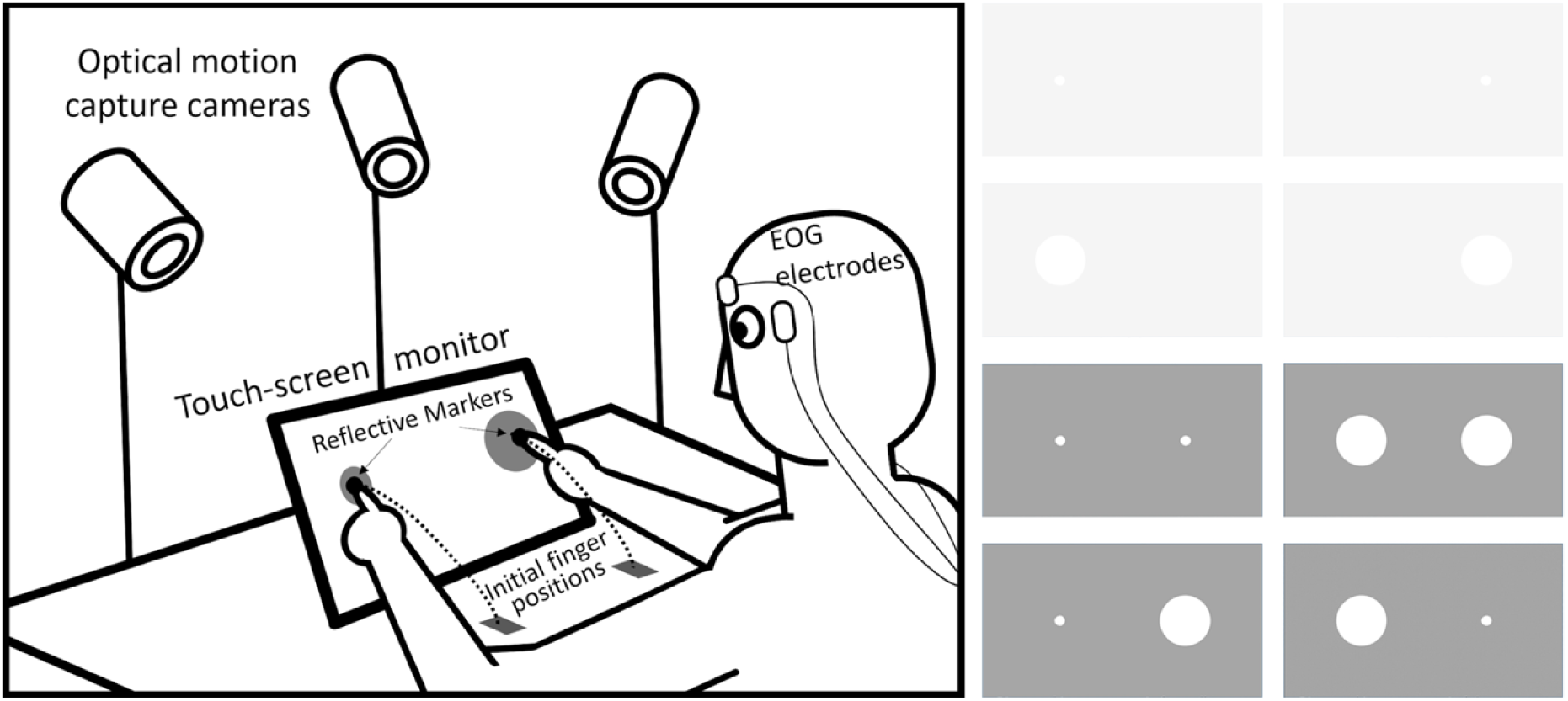
Experimental setup and target conditions. The four bimanual target conditions analyzed in the present study are shown in the lower half of the right-hand panel. Reproduced with permission from Sardar et al. [13].

For this analysis, we restricted the data to the strictly bimanual portion of each reach, defined as the interval from the onset of movement of the later-starting hand to the first target touch by either hand. After excluding four trials with target-touch asynchrony greater than 1.5 s and five trials with prolonged missing kinematic data, a total of 751 candidate bimanual trials were available. Fifteen trials did not contain a valid overlapping bimanual movement interval, leaving 736 trials for further analysis. Lateral gaze phases were defined as continuous periods in which gaze was directed toward either the left or right side. Only the portion of each gaze phase that overlapped with the strictly bimanual interval was included.

For each valid gaze phase, the average BTD slope was calculated using linear Deming regression. The initial gaze shift from central fixation toward either side was included in the descriptive count of gaze shifts, but it was not included in the consistency analysis because calculation of the change in BTD slope (δ) requires two consecutive lateral gaze phases.

Accordingly, only transitions from left to right gaze or from right to left gaze, named as “lateral gaze transitions”, for which valid BTD slopes could be calculated for both the preceding and following gaze phases were included in the consistency analysis. Of the 751 candidate bimanual trials, 15 did not contain a valid overlapping bimanual movement interval, leaving 736 trials eligible for the strict-bimanual analysis. Of these 736 trials, 555 contributed at least one eligible transition, yielding 794 transitions for the consistency analysis. Trials that did not contribute a transition were not treated as failed or invalid trials; rather, they did not contain an eligible gaze-transition event within the analysis window.

As described above, an increase in BTD slope indicates that the speed ratio shifted toward the right hand, whereas a decrease indicates that it shifted toward the left hand. A gaze shift was classified as consistent when the change in BTD slope favored the hand on the seen side, and as inconsistent when it favored the hand on the unseen side. Across participants, the mean proportion of consistent shifts was 76.1% ± 11.5% (mean ± SD; n = 19). This proportion exceeded chance, t(18) = 9.87, p <.001, 95% CI [70.6%, 81.7%]; the Wilcoxon signed-rank test also supported this result (p <.001). A transition-level mixed-effects logistic model estimated a consistency probability of 77.5% (odds ratio = 3.44, β = 1.24, SE = 0.12, p <.001). Thus, consistent shifts were the predominant eye–hand coordination pattern when the analysis was restricted specifically to the period in which both hands were moving.

To determine whether this finding depended on the particular BTD slope measure, we repeated the analysis using a simpler normalized relative-speed asymmetry measure,

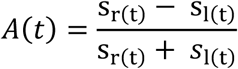

where positive values indicate relatively greater right-hand speed and negative values indicate relatively greater left-hand speed. For each gaze phase, *A*(*t*) was averaged over the strictly bimanual portion of that phase, and the difference in mean *A*(*t*) between two consecutive gaze phases was calculated. A rightward gaze shift was classified as consistent when this difference was positive, and a leftward gaze shift was classified as consistent when it was negative. Across the same 794 transitions from 19 participants, mean consistency was 82.2% ± 10.0%, t(18) = 14.07, p <.001, 95% CI [77.4%, 87.0%]. The mixed-effects logistic model estimated a consistency probability of 81.6% (odds ratio = 4.44, β = 1.49, SE = 0.14, p <.001). Thus, the association between gaze direction and relative hand dynamics was not dependent on the particular BTD slope formulation.

At the beginning of a bimanual reach, participants may look at both targets in sequence and initiate the corresponding hand movements in the same order. This initial tendency could bias the observed association between gaze direction and changes in relative hand speed. To assess this possibility, we repeated the analysis after excluding the first gaze transition in each trial. After excluding the first lateral gaze transition, 239 transitions from 17 participants remained. Mean consistency was 72.6% ± 17.8%, t(16) = 5.25, p <.001, 95% CI [63.5%, 81.8%]. The mixed-effects logistic model estimated a consistency probability of 68.4% (odds ratio = 2.16, β = 0.77, SE = 0.16, p <.001). Therefore, the predominance of consistent shifts cannot be attributed solely to the first lateral gaze transition within each trial.

Also, to examine whether this relationship depended on when gaze shifted during the reach, the strictly bimanual movement interval was divided into equal early, middle, and late thirds according to normalized elapsed time. Consistent shifts remained significantly more frequent than chance during all three stages. The early third included 183 transitions, with mean consistency of 75.6% ± 23.5%, t(18) = 4.75, p <.001, 95% CI [64.3%, 86.9%]. The middle third included 311 transitions, with mean consistency of 82.8% ± 14.4%, t(18) = 9.93, p <.001, 95% CI [75.9%, 89.8%]. The late third included 300 transitions, with mean consistency of 68.1% ± 18.1%, t(18) = 4.37, p <.001, 95% CI [59.4%, 76.8%]. All three stages included 19 participants. The corresponding mixed-effects logistic models estimated consistency probabilities of 75.4%, 85.9%, and 70.0%, respectively (early: odds ratio = 3.07, β = 1.12, SE = 0.17; middle: odds ratio = 6.08, β = 1.80, SE = 0.18; late: odds ratio = 2.33, β = 0.85, SE = 0.18; all p <.001). This indicates that the relationship was present throughout the bimanual reach rather than being restricted to a specific sub-period, either the beginning or the end, of the movement.

Taken together, these results suggest that a consistent shift, wherein the speed ratio shifts toward the hand on the seen side, is the predominant eye–hand coordination pattern during bimanual reaching movements.

## DISCUSSION

We proposed a new analysis of hand-eye coordination during bimanual reaching that simplifies a pair of three-dimensional trajectories into two-dimensional trajectories of traveled distance, referred to as BTD. Using this approach, a succinct framework for examining the competitive dynamics of two hands was established, focusing on changes in the relative speeds between the two hands associated with gaze shifts. We then analyzed the consistency between the change in gaze direction and that in BTD slope and demonstrated that the asynchronous bimanual dynamics strongly correlate with gaze-shifting patterns. Importantly, this relationship remained when the analysis was restricted to the strictly bimanual portion of the movement and was also reproduced using an alternative normalized relative-speed measure. The relationship was observed throughout the early, middle, and late stages of the bimanual movement and remained significant after excluding the first lateral gaze transition within each trial. Together, these findings suggest that the relationship between gaze direction and relative hand dynamics is a robust feature of visually guided bimanual reaching rather than being restricted to a particular portion of the movement or dependent on the specific BTD formulation.

### Comparison with previous studies

While few studies have previously examined both hand and eye movements during bimanual reaching tasks, there is good alignment between our broader observations and these previous studies. The number of gaze shifts observed during each trial in our study (median = 2) was comparable with that reported by Riek et al. [9] and Bruyn and Mason [5].

Also comparable with earlier studies, we found some trials were completed without detected lateral gaze phases; however, the proportion we observed was substantially smaller (2.7%) compared to the 38% reported by Honda [15] and the 20% and 5% found respectively for congruent and incongruent bimanual reaches reported by Bruyn and Mason [5]. Such differences are likely due to task differences and the related accuracy demands. The lateral bias observed in gaze behavior in our experiment also appears consistent with previous studies. We direct readers to our previous paper that focused on more conventional analyses of the same data set [13] (see section “An initial consideration of eye movement data”), which confirmed the previously observed tendency for gaze to be directed more toward the dominant side [14, 15] and/or to direct gaze more toward targets requiring greater reaching precision [9].

In comparison to previous studies, our model provides a finer-grained understanding of how visual guidance is associated with bimanual activities. Studies suggest that gaze shifts during bimanual reaching movements facilitate visual guidance of the seen hand, thereby assigning control priority to it compared to the unseen hand [2, 4–6]. Although such gaze shifts may not be a necessary condition for bimanual asynchrony, as demonstrated when gaze is restricted [5, 6, 11, 12, 16], our data, along with other studies [4, 9, 17], show that gaze shifting is a strongly preferred strategy under natural viewing conditions. The present study consolidates these observations by demonstrating a systematic association between gaze shifts and changes in the relative dynamics of the two hands.

Existing models explaining how bimanual asynchrony arises refer to different phases of bimanual movement, either during reaching [9, 10] or reach-to-grasp activities [4, 5]. The related classification of phases (or strategies) using bespoke criteria of hand kinematics includes the first minimum of speed [9], the wrist being stationary [4], and differences in hand arrival time [10]. These investigations, focused on interpreting bimanual asynchrony based on these classifications, have contributed to our understanding of bimanual behavior but have not provided a more general understanding of the relationship between gaze shifts and bimanual coordination. These previous studies, together with our own earlier analysis [13], were therefore limited in their ability to explain how gaze shifts relate to the changing coordination of the two hands throughout the movement.

Our model provides evidence of a systematic correlation between gaze and hand movements throughout the bimanual movement. Importantly, this does not conflict with existing models but extends them by examining relative hand dynamics at a finer temporal scale. Previous studies have argued that asynchrony becomes particularly evident toward the end of the movement, where the leading hand may remain stationary while waiting for the following hand. This virtually serial part of the movement has been referred to as the completion component [4], hover phase [9], or sequential placement [10]. Our primary analysis specifically excluded this terminal unimanual period by restricting the analysis to the interval in which both hands were moving. Nevertheless, consistent gaze–hand relationships were observed during the early, middle, and late thirds of this strictly bimanual interval.

Thus, the observed relationship cannot be explained solely by the terminal phase in which one hand has already completed its movement. Rather, subtle changes in relative hand dynamics associated with gaze direction are evident throughout the simultaneously executed bimanual movement.

This finding also does not conflict with the possibility that gaze may serve different functions at different stages of reaching. Gaze directed toward a target early in the movement may primarily support target localization and directing the hand toward the target region, whereas gaze later in the movement may increasingly support online guidance and accurate contact [18–20]. Despite these potentially different functional roles, consistent gaze–hand relationships were significantly more frequent than chance in each third of the movement.

In addition, what makes our approach different from existing models is that it does not attempt to define movement phases using arbitrary hand-kinematic criteria. Rather, the principal segmentation is based on gaze behavior, which provides naturally occurring periods of leftward and rightward visual orientation. We then focus on the competitive bimanual dynamics represented as changes in the BTD slope across these gaze phases. Importantly, the present analysis was restricted to the strictly bimanual part of the movement, excluding periods before both hands were moving and after either hand had reached the target.

Therefore, the reported relationship specifically reflects changes in relative hand dynamics while both component movements were ongoing.

### Interpretation of ***δ***

Our model is based on a new variable δ, which is the difference in average speed ratio across neighboring gaze phases. Although this is a new variable devised based on empirical observation, its meaning can be understood by looking at its mathematical structure. Note that the speed ratio σ, i.e., the slope of the BTD trajectory, is defined as:

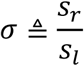

where *s_r_* and *s_l_* are the speeds of the right and left hands, respectively. The differential of this ratio is

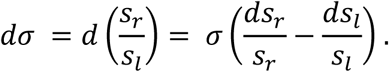

This differential describes local changes in the speed ratio, whereas δ summarizes the difference between fitted BTD slopes across consecutive gaze phases. Thus, a positive δ indicates that the speed ratio shifted toward the right hand, whereas a negative δ indicates that the relative speed ratio shifted toward the left hand. In the present analysis, only the sign of δ was considered. Therefore, δ provides a phase-level indication of the direction of change in the relative dynamics of the two hands.

An obvious question is why the analysis is based on δ, an approximation obtained by comparing the average slopes of BTD segments, instead of using an instantaneous measure of the difference. We compared phase-level BTD orientations to summarize changes across gaze phases without assuming that gaze shifts and changes in hand dynamics occurred at exactly the same instant. It is unlikely that a gaze shift and a corresponding change in hand dynamics are perfectly synchronized; for example, changes in relative hand dynamics may follow the gaze shift because of visuomotor delay, or may begin beforehand because of the complex bidirectional relationship between eye and hand movement [21–23]. While our analysis could have incorporated a free parameter to estimate the delay between gaze and hand movement, this approach was avoided to prevent potential overfitting.

The complementary analysis using normalized relative-speed asymmetry, A(t), reproduced the association between gaze-shift direction and changes in relative hand dynamics, supporting the conclusion that this association is not specific to the BTD formulation.

Indeed, the proportion of gaze-consistent transitions was higher for the A(t)-based analysis than for the δ-based analysis (82.2% versus 76.1%). Nevertheless, we retained δ as the primary measure because it directly connects the geometric representation of bimanual movement to the comparison of successive gaze phases. By estimating a representative BTD orientation within each phase, δ characterizes how the overall trajectory direction changes across a gaze transition, providing a coherent link between visualization and quantitative analysis. The higher consistency proportion obtained with A(t) provides valuable corroborating evidence, but does not by itself establish that phase-averaged speed asymmetry is a more appropriate primary measure for this framework.

### Implications and Limitations

One possible functional interpretation for this phenomenon is that directing visual attention toward one side facilitates visual guidance of the movement on that side. As in other examples of eye-hand coordination [24–26], the observed association may therefore reflect a dynamic, task-dependent allocation of visual attention between the two component movements.

More generally, bimanual synchrony and asynchrony should be considered in relation to the specific demands of the task rather than as fixed properties of bimanual control. The strong overall synchrony observed here coexisted with small within-movement changes in the relative progression of the two hands. Such flexibility is consistent with the view that bimanual coordination reflects a balance between coupling the two component movements and satisfying the task-specific demands imposed on each limb. In the present task, visual information could normally be acquired from only one side at a time, providing a plausible task constraint under which this balance is continuously adjusted as the movement unfolds.

A primary limitation of our model is that it remains largely descriptive rather than generative. While it satisfactorily describes patterns of eye-hand bimanual coordination, it does not offer a systematic method for simulating these patterns. Additionally, this is a highly simplified model, reducing the total seven degrees of freedom in the captured bimanual eye-hand movements (three-dimensional for each hand and one-dimensional lateral gaze behavior) to an analysis of the association between gaze direction and phase-to-phase changes in relative hand speed. Although the succinct correlations reported in our study might have only become observable through this simplification procedure, it inevitably overlooks the complexity of the full coordination dynamics. More generally, these limitations underscore the need for further development toward a fully generative model. Future research should focus on integrating generative elements into the model to achieve a comprehensive framework for eye-hand coordination during bimanual tasks by incorporating computational frameworks such as optimal feedback control [27].

Another limitation of our model is that it does not explain causality in eye-hand coordination. This means that it reports the correlation and remains agnostic about whether gaze shifts influence hand asynchrony, whether changes in hand dynamics influence gaze allocation, or whether both are consequences of a common control process. For the same reason, we remain uncertain whether the gaze and hand patterns are determined in an offline, feedforward manner or whether they result from online feedback corrections. To address this, future work could manipulate gaze behavior using fixation or visual or auditory cues, or mechanically perturb ongoing hand movements. Such approaches would help determine the directional influence between gaze shifts and hand asynchrony, providing a deeper understanding of the underlying mechanisms in bimanual coordination.

Finally, EOG provides a robust measure of horizontal gaze direction but does not identify the precise location being fixated. Consequently, the present analysis can distinguish whether gaze was directed toward the left or right side but cannot determine whether participants were looking specifically at the target, their moving finger, or another location on the same side. Future studies using calibrated video-based eye tracking could distinguish these possibilities and establish how different fixation locations relate to the evolving dynamics of the two hands.

## METHODS

Nineteen healthy individuals, aged 18 to 23 years, took part in the experiment. Ten of these participants were right-handed, while the remainder were left-handed, as confirmed by the Edinburgh Handedness Inventory [28]. All participants had normal vision and were free from any known neurological or musculoskeletal disorders. The project was reviewed and approved by the University of Birmingham’s Science, Technology, Engineering and Mathematics Ethical Review Committee, and all participants provided their informed consent before participating.

### Motion and Eye Tracking

Participants’ bimanual movements were measured using a commercial motion capture system (ProReflex, Qualisys AB, Gothenburg, Sweden). This system comprised three cameras positioned around the workspace, which tracked the three-dimensional positions of 5 mm reflective markers attached to the nails of both index fingers at a sampling rate of 200 Hz.

In synchrony with the motion tracking, horizontal eye movements were recorded using electro-oculography (EOG). Self-adhesive surface electrodes were placed on the canthi of both eyes, with a ground electrode attached to the glabella. The EOG signals were sampled at 2 kHz and amplified (2 K) and band-pass filtered (0.1–30 Hz) via an AC preamplifier (Grass Instruments LP122).

### Experimental Procedure

Participants were seated in front of a 23-inch LCD touchscreen HD monitor (Dell S2340T), which was placed on a table tilted backwards at a 27° angle from vertical. To begin each trial, participants were instructed to position their left and right index fingers on two marked starting points on the table, spaced 25 cm apart. From the midpoint of these two starting points, the center of the touchscreen monitor was located 35 cm away horizontally and 16.5 cm away vertically, resulting in a straight-line distance of 38.7 cm from the midpoint, with a slope of 25.2° from the horizontal plane.

While maintaining their index fingers on the starting positions, participants were instructed to focus on a fixation cross displayed at the center of the screen for a randomly selected duration per trial, ranging from 1 to 3 seconds. Subsequently, the fixation cross disappeared, and targets appeared. Participants were directed to make reaching movements toward the targets as fast and as accurately as possible. The targets could appear on the left, right, or both sides of the display, prompting participants to perform left, right, or bimanual movements accordingly. The collected unimanual and bimanual data were previously used in the authors’ paper on the same dataset [13]. Since this study is specifically focused on eye-movement behavior during bimanual reaching, only bimanual trials were used in the present analysis.

Target sizes varied from small (diameter = 2 cm) to large (diameter = 10 cm). The distance between the centers of the left and right targets remained constant at 35 cm (approximately 14°), and the two centers always aligned horizontally. However, their midpoint was randomly displaced horizontally and vertically from the screen center, with a maximum allowed displacement of 5.29 cm. This was to encourage participants to rely more on visual information rather than memory. Upon touching the targets, participants withdrew their hands from the screen, and the touch locations were marked as small red dots for 1 second.

Since each side could feature a large, small, or no target, the original experiment consisted of eight different reaching conditions: four unimanual and four bimanual. The four bimanual conditions comprised small targets on both sides, large targets on both sides, a small target on the left and large target on the right, and a large target on the left and small target on the right. Following a practice session covering all eight conditions once each, participants completed ten blocks of eight trials, with conditions randomly permuted within each block.

## Data Processing and Analysis

Motion and EOG data were processed in MATLAB (The MathWorks Inc., Natick, MA, USA). Three-dimensional fingertip trajectories were low-pass filtered using a fourth-order Butterworth filter with a cutoff frequency of 20 Hz. Hand speed was calculated as the magnitude of the first derivative of the filtered trajectories. Movement onset was identified as the start of a monotonic increase in speed that subsequently exceeded 40 cm/s.

EOG signals were low-pass filtered using a fourth-order Butterworth filter with a cutoff frequency of 30 Hz and baseline-corrected by subtracting the mean of the first 0.5 s of each trial during central fixation. Lateral saccade onset was identified as the start of a monotonic increase in absolute EOG amplitude that subsequently exceeded 0.15 V; signal polarity distinguished leftward from rightward gaze. A gaze phase comprised a continuous period of gaze toward the same lateral side. Unclassified gaps of up to 20 EOG samples (10 ms) were bridged when flanked by gaze toward the same side.

Consistency analyses were restricted to the interval from movement onset of the later-starting hand to the first target touch by either hand. Only the portion of each gaze phase overlapping this interval was retained. Eligible transitions were left-to-right or right-to-left shifts with valid estimates for both adjacent phases. The initial shift from central fixation to either side was counted in the descriptive gaze-shift analysis but excluded from consistency analyses.

Traveled distance was calculated as the cumulative sum of Euclidean distances between successive filtered fingertip positions. BTD trajectories plotted right-hand against left-hand traveled distance. For each gaze phase, a representative BTD slope, σ̅*_p_*, was estimated using linear Deming regression with an error-variance ratio of 1 and a freely estimated intercept. The change between consecutive phases was defined as *δ* = σ̅*_p_*_+1_ − σ̅*_p_*. A transition was classified as consistent when *δ* > 0 for a left-to-right shift or *δ* < 0 for a right-to-left shift; opposite sign combinations were classified as inconsistent.

Robustness was assessed using normalized relative-speed asymmetry, *A*(*t*) = [*s_r_*(*t*) − *s_l_*(*t*)]/[*s_r_*(*t*) + *s_l_*(*t*)], where *s_r_*(*t*) and *s_l_*(*t*) are instantaneous right-and left-hand speeds. This measure was averaged over the retained portion of each gaze phase, and changes between consecutive phase means were classified using the same directional criterion. Additional analyses repeated the BTD consistency analysis after excluding the first eligible lateral gaze transition in each trial, and separately for transitions occurring within the early, middle, and late thirds of the time-normalized bimanual interval.

## Statistical Analysis

The number of gaze shifts per trial, including zero-shift trials, was analyzed using a Poisson mixed-effects model with target-size combination as a fixed effect and participant as a random intercept; Bonferroni adjustment was applied to the three comparisons of the small-small reference condition with each other target-size combination. For each participant, the proportion of eligible gaze transitions classified as consistent was calculated. Participant-level consistency proportions were compared with the chance level of 0.5 using a one-sample t-test, with 95% confidence intervals reported. A Wilcoxon signed-rank test against 0.5 was additionally performed as a non-parametric sensitivity analysis.

Because individual participants and trials could contribute multiple gaze transitions, the binary transition-level consistency outcome was also analyzed using a generalized linear mixed-effects model with a binomial distribution and logit link: Consistent ∼ 1 + (1 | Participant) + (1 | Participant:Trial).

Random intercepts were therefore included for participant and trial nested within participant. Models were fitted in MATLAB using fitglme, with Laplace approximation for the Poisson count model and maximum pseudo-likelihood for the binomial consistency models. Fixed-effect estimates are reported on the log-odds scale together with standard errors, odds ratios, consistency probabilities evaluated with the random effects set to zero, and p-values.

The same inferential procedure was applied to the alternative relative-speed measure and to the sensitivity analysis excluding the first eligible gaze transition. For the movement-stage analysis, consistency was examined separately during the early, middle, and late thirds of the strictly bimanual interval. These stage-specific analyses were used to determine whether the gaze–hand relationship was present throughout the movement rather than to test statistical differences between movement stages.

## ACKNOWLEDGEMENTS

We thank Jon Allsop for help with the experimental setup.

## AUTHOR CONTRIBUTIONS

S.-H.Y. contributed to conceptualization, methodology, formal analysis, writing of the original draft, review and editing of the manuscript, and supervision. S.D.S. contributed to methodology, data curation, and review and editing of the manuscript. C.M. contributed to review and editing of the manuscript. D.P. contributed to conceptualization, supervision, methodology, and review and editing of the manuscript. All authors reviewed and approved the final manuscript.

## DATA AVAILABILITY

The datasets generated during and/or analyzed during the current study are available from the corresponding author on reasonable request.

## ADDITIONAL INFORMATION

Competing interests: The authors declare no competing interests.

Correspondence and requests for materials should be addressed to S.-H. Yeo.

Use of artificial intelligence tools: Generative AI tools were used solely for language proofreading and minor editorial assistance during manuscript preparation. The authors reviewed and approved all changes and take full responsibility for the final content of the manuscript.

